## Supplementary Figures and Tables for "*Pseudomonas aeruginosa* PA14 produces R-bodies, extendable protein polymers with roles in host colonization and virulence"

### Suppl. Figure 1

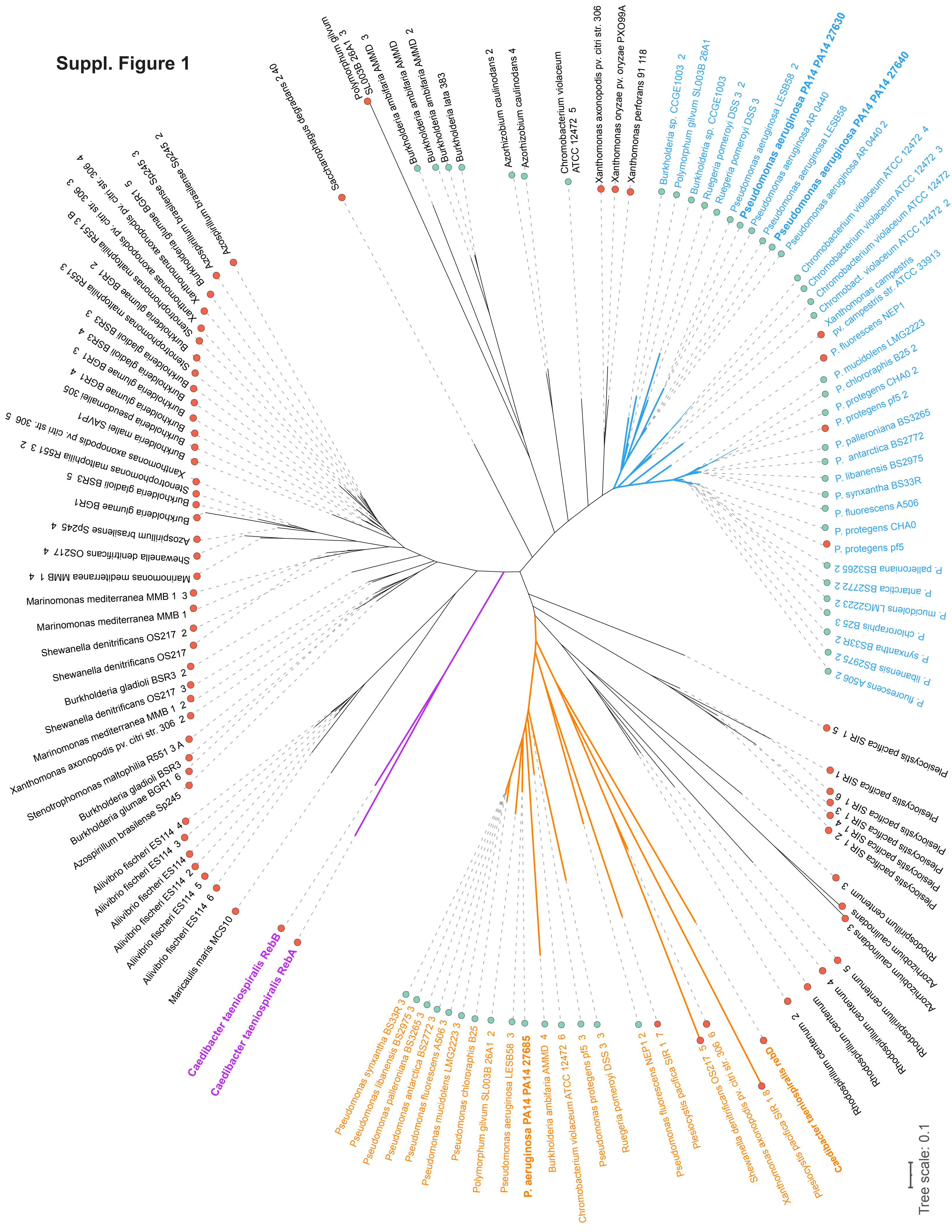

**Supplementary Figure 1.** Phylogenetic tree of proteins from 38 bacterial strains that are homologous to *C. taeniospiralis* RebB. The bacterial strains included are those with complete genomes from Figure 6 of Raymann *et al*. (1) plus 14 representative pseudomonads. *C. taeniospiralis* RebA and RebB are shown in purple. The cluster containing *C. taeniospiralis* RebD and its *Pseudomonas* homologs is shown in orange. We have assigned the designation "RebP" to Reb homologs that cluster with *P. aeruginosa* PA14_27630 and PA14_27640 (shown in blue). Green circles at the end of each branch indicate the presence of FecI2 homologs in the respective strains, while red circles indicate its absence.

**Suppl. Figure 2**

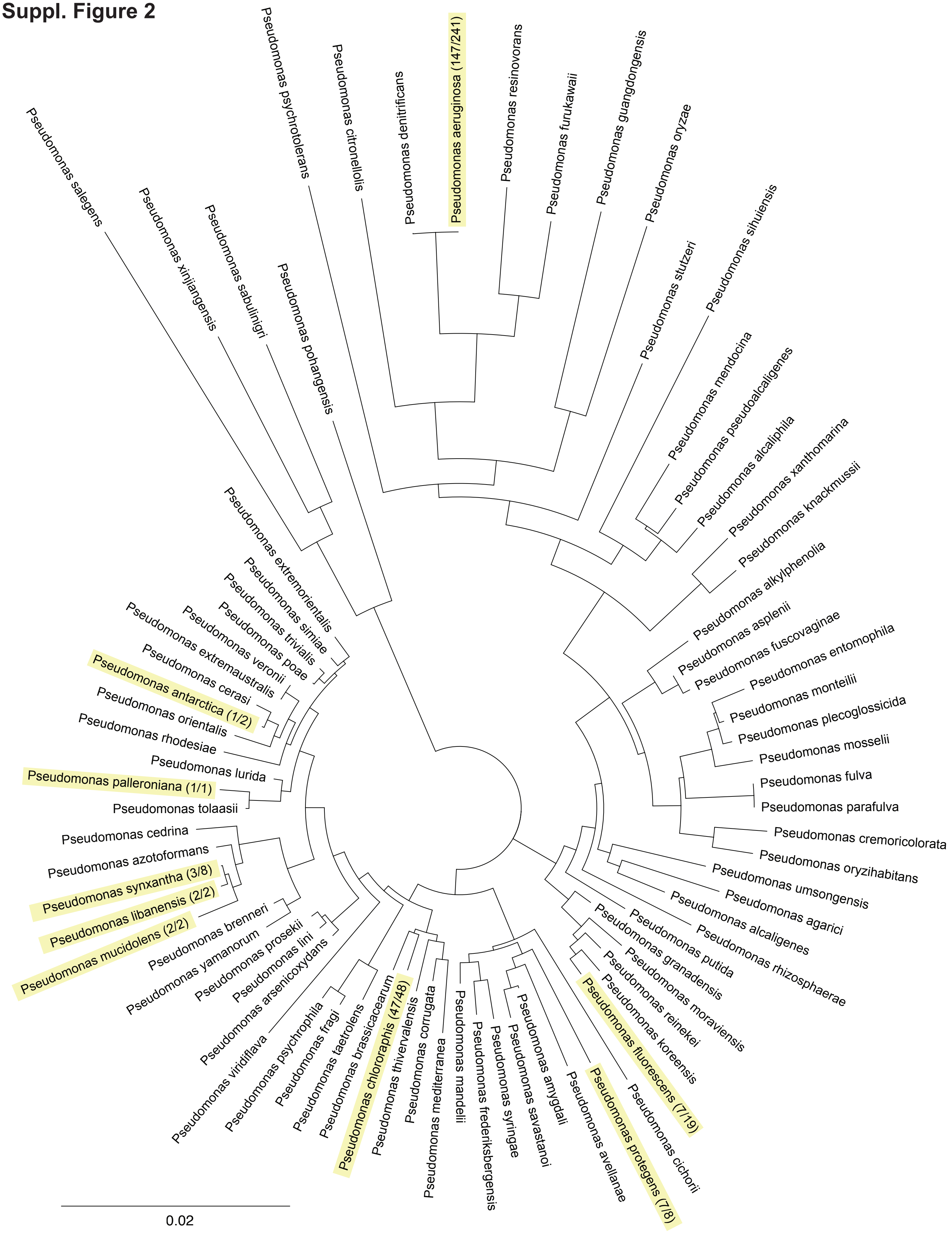

**Supplementary Figure 2.** 16S rRNA-based phylogenetic tree showing all *Pseudomonas* species with complete genomes available in the Pseudomonas Genome Database. Species with at least one strain containing one or more *rebP1* homolog(s) are highlighted in yellow. Indicated in parentheses are the number of strains with *rebP1* homologs out of the total number of strains with complete genomes for a respective species.

**Suppl. Figure 3**

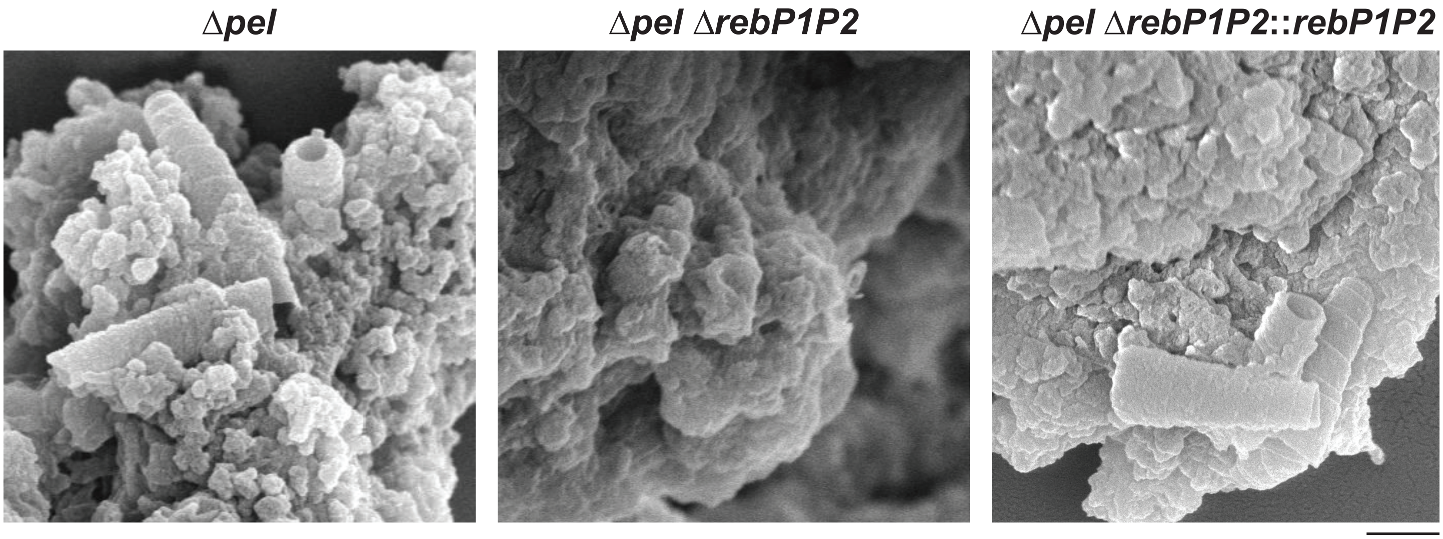


**Supplementary Figure 3.** SEM images of SDS-insoluble fractions prepared from biofilms of strains with the indicated mutations. The Δ*pel* background was used because this parent strain is more amenable to disruption. Scale bar is 500 nm. The images are representative of 16 fields of view captured for each strain. For Δ*pel* and Δ*pel*Δ*rebP1P2*::*rebP1P2* samples, every field of view captured had an R-body present whereas none were detected in the Δ*pel*Δ*rebP1P2* sample. On average, 10.8 R-bodies were seen in each field of view (range from 1-40 R-bodies) for Δ*pel* and 9.6 R-bodies with a range of 1-30 R-bodies in Δ*pel*Δ*rebP1P2*::*rebP1P2*.

**Suppl. Figure 4**

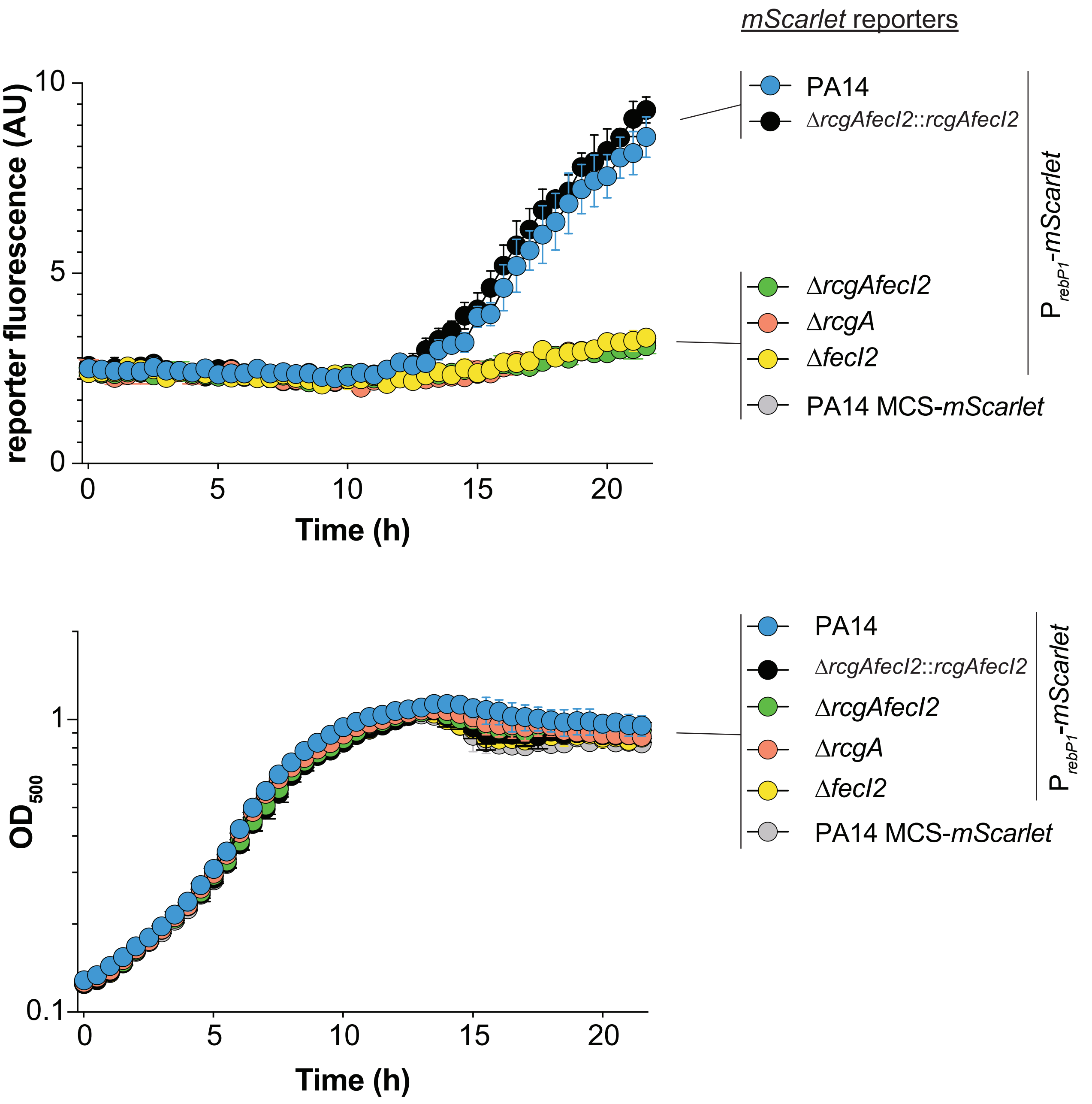


**Supplementary Figure 4.** Expression of mScarlet by the indicated genotypes containing the $P_{rebP1}$ or promoterless transcriptional reporter in shaken 1% tryptone liquid cultures grown at 25 °C. Corresponding growth curves are shown in the bottom panel.

**Suppl. Figure 5**

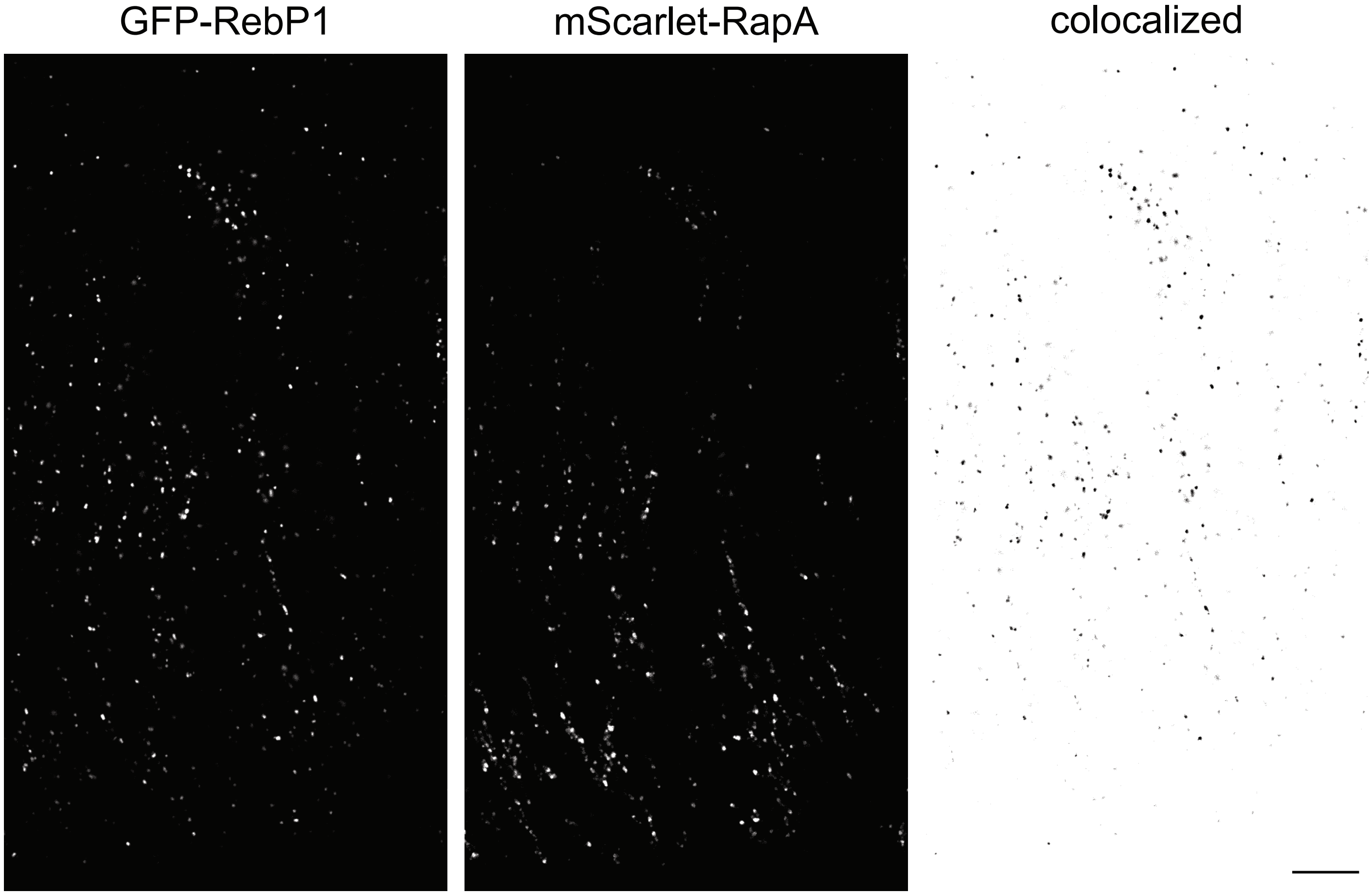


**Supplementary Figure 5.** Colocalization of GFP-RebP1 and mScarlet-RapA. Representative confocal image of a thin section prepared from three-day-old biofilms of the dual-labeled strain shown in **Figure 4e**. To visualize the degree of colocalization, we generated a binary mask of GFP-RebP1 in ImageJ. The mask was then applied (ImageJ image calculator, Subtract) to the mScarlet-RapA image leaving only the colocalized fraction of pixels as shown on the right. Scale bar is 10 µm.

#### Supplementary Table 1: Strains

| Strain | Number | Description | Source | Appears in figure |
|---|---|---|---|---|
| *Pseudomonas aeruginosa* | | | | |
| UCBPP-PA14 | | Clinical isolate UCBPP-PA14. | (2) | 2b-e, 3e-f, 4c-d, Table S2 |
| PA14 Δ*pel* | LD82 | PA14 with deletion in *PA14_24490 (pelB)-24560 (pelG).* | (3) | 2b-e, S3, Table S2 |
| PA14 Δpel ΔrebP1P2 | LD2720 | PA14 with deletion in *PA14_24490-24560* and *PA14_27640 (rebP1)-27640 (rebP2).* Made by mating pLD2934 into LD2720. | This study | S3 |
| PA14 Δpel ΔrebP1P2:: *rebP1P2* | LD3026 | PA14 Δ*pelB-G* Δ*PA14_27640 (rebP1)-27630 (rebP2)* with wild-type *PA14_27630-27640* complemented back into the site of deletion. Made by mating pLD3016 into LD2936. | This study | S3 |
| PA14 $P_{rebP1}$-*mScarlet* | LD3224 | PA14 with $P_{rebP1}$-*mScarlet* inserted at the *attB* site using pLD3210. | This study | 3a, 4a-b |
| PA14 Δ*rcgAfecI2* | LD3137 | PA14 with deletion in *PA14_27690 (fecI2) -27700 (rcgA)*. Made by mating pLD3136 into UCBPP-PA14. | This study | - |
| PA14 Δ*fecI2* | LD3144 | PA14 with deletion in *PA14_27690 (fecI2)*. Made by mating pLD3142 into UCBPP-PA14. | This study | - |
| PA14 Δ*rcgA* | LD3145 | PA14 with deletion in *PA14_27700 (rcgA)*. Made by mating pLD3143 into UCBPP-PA14. | This study | - |
| PA14 Δ*rcgAfecI2*:: *rcgA-fecI2* | LD3180 | PA14 Δ*PA14_27700 (rcgA)-27690 (fecI2)* strain with wild-type *PA14_27700-27690* complemented back into the site of deletion. Made by mating pLD3179 into LD3137. | This study | - |
| PA14 Δ*rcgAfecI2* $P_{rebP1}$-*mScarlet* | LD3782 | PA14 Δ*PA14_27690 (rcgA)-27700 (fecI2)* with $P_{rebP1}$-*mScarlet* inserted at the *attB* site using pLD3210. | This study | 3a |
| PA14 Δ*fecI2* $P_{rebP1}$-*mScarlet* | LD3780 | PA14 Δ*PA14_27690 (fecI2)* with $P_{rebP1}$-*mScarlet* inserted at the *attB* site using pLD3210. | This study | 3a |
| PA14 Δ*rcgA* $P_{rebP1}$-*mScarlet* | LD3781 | PA14 Δ*PA14_27700 (rcgA)* with $P_{rebP1}$-*mScarlet* inserted at the *attB* site using pLD3210. | This study | 3a |

| | | | | |
|---|---|---|---|---|
| PA14 Δ*rcgAfecI2*:: *rcgAfecI2* P*rebP1*-*mScarlet* | LD3783 | PA14 Δ*PA14_27700 (rcgA)-27690 (fecI2)* strain with wild-type *PA14_27700-27690* complemented back into the site of deletion. The strain also has P*rebP1*-*mScarlet* inserted at the *attB* site using pLD3210. | This study | 3a |
| PA14 MCS-*mScarlet* | LD3294 | PA14 without a promoter driving *mScarlet* expression inserted at the *attB* site using pLD3208. | This study | 3a, 4a-b |
| PA14 P*rebP1*-*mScarlet* P*PA1/04/03*-*gfp* | LD3657 | PA14 with P*rebP1*-*mScarlet* inserted at the *attB* site and P*PA1/04/03*-*gfp* inserted at the *glmS* site by mating pLD3655 into LD3224. | This study | 3b-d |
| PA14 Δ*rcgAfecI2* P*rebP1*-*mScarlet* P*PA1/04/03*-*gfp* | LD3786 | PA14 Δ*PA14_27700 (rcgA)-27690 (fecI2)* with P*rebP1*-*mScarlet* inserted at the *attB* site using pLD3210 and P*PA1/04/03*-*gfp* inserted at the *glmS* site by mating pLD3655 into LD3782. | This study | 3c-d |
| PA14 Δ*fecI2* P*rebP1*-*mScarlet* P*PA1/04/03*-*gfp* | LD3784 | PA14 Δ*PA14_27690 (fecI2)*with P*rebP1*-*mScarlet* inserted at the *attB* site and P*PA1/04/03*-*gfp* inserted at the *glmS* site by mating pLD3655 into LD3780. | This study | 3c-d |
| PA14 Δ*rcgA* P*rebP1*-*mScarlet* P*PA1/04/03*-*gfp* | LD3785 | PA14 Δ*PA14_27700 (rcgA)* with P*rebP1*-*mScarlet* inserted at the *attB* site and P*PA1/04/03*-*gfp* inserted at the *glmS* site by mating pLD3655 into LD3781. | This study | 3c-d |
| PA14 Δ*rcgAfecI2*:: *rcgA-fecI2* P*rebP1*-*mScarlet* P*PA1/04/03*-*gfp* | LD3787 | PA14 Δ*PA14_27700 (rcgA)-27690 (fecI2)* strain with wild-type *PA14_27700-27690* complemented back into the site of deletion. The strain also has P*rebP1*-*mScarlet* inserted at the *attB* site and P*PA1/04/03*-*gfp* inserted at the *glmS* site by mating pLD3655 into LD3783. | This study | 3c-d |
| PA14 Δ*rebP1P2* | LD2935 | PA14 with deletion in *PA14_27640 (rebP1)-27630 (rebP2)*. Made by mating pLD2934 into UCBPP-PA14. | This study | 3e-f, 4c-f |
| PA14 P*PA1/04/03*-*mScarlet* | LD3765 | PA14 constitutively expressing mScarlet. Made by mating pLD3433 into UCBPP-PA14. | This study | 3f |
| PA14 Δ*rebP1P2* P*PA1/04/03*-*mScarlet* | LD3766 | PA14 Δ*PA14_27640 (rebP2)-27630 (rebP2)* constitutively expressing mScarlet. Made by mating pLD3433 into LD2935. | This study | 3f |
| PA14 P*PA1/04/03*-*mScarlet* | LD3295 | PA14 with *lac*-derived constitutive *PA1/04/03* promoter driving mScarlet expression inserted at the *attB* site using pLD3293. | This study | 4a-b |

| | | | | |
|---|---|---|---|---|
| PA14 Δ*rebP1P2*::*rebP1P2* | LD3025 | PA14 Δ*PA14_27640 (rebP1)-27630 (rebP2)* with wild-type *PA14_27640-27630* complemented back into the site of deletion. Made by mating pLD3016 into LD2935. | This study | 4c-d |
| PA14 *gacA*::*Tn* | LD1560 | *MAR2xT7* transposon insertion into *PA14_30650 (gacA)*. | (4) | 4c-d |
| *Arabidopsis thaliana* | | | | |
| *Col-0* | | | J. Reed, UNC-Chapel Hill | 4a,c |
| *Caenorhabditis elegans* | | | | |
| *unc-44(e362)* | LD3326 | | (5) M. Chalfie, Columbia University | 4b,d |
| *Escherichia coli* | | | | |
| UQ950 | LD44 | *E. coli* DH5α λ(*pir*) strain for cloning; F- Δ(*argF-lac*) 169φ80d*lacZ*58(ΔM15) *glnV44*(AS) *rfbD1 gyrA96*(NalR) *recA1 endA1 spoT thi-1 hsdR17 deoR λpir+* | D. Lies, Caltech | |
| BW29427 | LD661 | Donor strain for biparental conjugation; *thrB1004 pro thi rpsL hsdS lacZ* ΔM15RP4-1360 Δ(*araBAD*)*567* Δ*dapA1341*::[*erm pir*(wt)] | W. Metcalf, University of Illinois | |
| S17-1 | LD2901 | StrR, TpR, F-RP4-2-*Tc::Mu aphA::Tn7 recA λpir* lysogen | (6) | |
| ß2155 | LD69 | Helper strain. *thrB1004 pro thi strA hsdsS lacZ*ΔM15 (F′ *lacZ*ΔM15 *lacI*$^{q}$ *traD36 proA*$^{+}$ *proB*$^{+}$) Δ*dapA*::*erm* (Erm$^{r}$) *pir*::RP4 [::kan (Km$^{r}$) from SM10] | (7) | |
| *Saccharomyces cerevisiae* | | | | |
| InvSc1 | LD676 | *MATa/MATα leu2/leu2 trp1-289/trp1-289 ura3-52/ ura3-52 his3-Δ1/his3-Δ1* | Invitrogen | |

**Supplementary Table 2: Primers**

| Primer number | Sequence | Used for plasmid |
|---|---|---|
| LD2609 | acgtacgtctcgagtctagatttaagaaggagatatacatatgagtaaaggagaagc | pLD3208 |
| LD2635 | actgactggagctcataaaacgaaaggcccagtctttcg | |
| LD2113 | acgtacgtacACTAGTccagatcctgcagaacgtc | pLD3210 |
| LD2114 | acgtacgtacGAATTCgatgtgactccctgtgagtgaa | |
| LD1087 | AGGGCCAATCGATAGAGTTT | |
| LD1088 | TCTTCGTGATCTGAAGCCATT | |
| LD3139 | acgtacgtacgcatgctg AGTAAAGGAGAAGAACTTTTCACTGG | pLD3655 |
| LD3141 | acgtacgtacgctagc GGCGGATTTGTCCTACTCAG | |
| LD2731 | acgtacgtacACTAGTtatttagaaaaataaacaaataggggttccgc | pLD3293 |
| LD2732 | acgtacgtacGAATTCgcttaatttctcctctttaattctagatgtgtg | |
| LD2507 | ggaattgtgagcggataacaatttcacacaggaaacagctCTACGATTGGGTGTCCTTGC | pLD3136 (LD2507-2510),<br><br>pLD1853 (LD2507 and LD2510) |
| LD2508 | TGTTCACGCCACTACACCCGCGACGTTTCACAAGACAGA | |
| LD2509 | TCTGTCTTGTGAAACGTCGCGGGTGTAGTGGCGTGAACA | |
| LD2510 | aggcaaattctgttttatcagaccgcttctgcgttctgatAGCGCTTCGACGAACAAC | |
| LD2507 | ggaattgtgagcggataacaatttcacacaggaaacagctCTACGATTGGGTGTCCTTGC | pLD3142 |
| LD2521 | TATGGATCGTCCGAATCAGCGCGACGTTTCACAAGACAGA | |
| LD2522 | TCTGTCTTGTGAAACGTCGCGCTGATTCGGACGATCCATA | |
| LD2523 | aggcaaattctgttttatcagaccgcttctgcgttctgatCTCGCTACCCTTTCCGAATA | |
| LD2525 | ggaattgtgagcggataacaatttcacacaggaaacagctACATGTCTGGGCACTCCTG | pLD3143 |
| LD2526 | CACGCCACTACACCCTGCCTGTCACCCAGGTAACAGCC | |
| LD2527 | GGCTGTTACCTGGGTGACAGGCAGGGTGTAGTGGCGTG | |
| LD2510 | aggcaaattctgttttatcagaccgcttctgcgttctgat AGCGCTTCGACGAACAAC | |
| LD2191 | ggaattgtgagcggataacaatttcacacaggaaacagctCGCGCGCAACTCTTCTAT | pLD2934 (LD2191-LD2194),<br><br>pLD3016 (LD2191 and LD2194) |
| LD2192 | gagttttccgaccgcagtcCTGCTGACGGTGCTCAAAG | |
| LD2193 | ctttgagcaccgtcagcagGACTGCGGTCGGAAAACTC | |
| LD2194 | aggcaaattctgttttatcagaccgcttctgcgttctgatGAAATATCGGACAGCGATGC | |

| | | |
|---|---|---|
| LD2507 | ggaattgtgagcggataacaatttcacacaggaaacagct CTACGATTGGGTGTCCTTGC | pLD3136 (LD2507-LD2509), pLD3179 (LD2507 and 2509) |
| LD2508 | TGTTCACGCCACTACACCCGCGACGTTTCACAAGACAGA | |
| LD2509 | TCTGTCTTGTGAAACGTCGCGGGTGTAGTGGCGTGAACA | |
| LD2510 | aggcaaattctgttttatcagaccgcttctgcgttctgat AGCGCTTCGACGAACAAC | |
| LD2811 | acgtacgtGCTGAGC AGTAAAGGAGAAGCTGTGATTAAAG | pLD3433 |
| LD2812 | acgtacgtGCT GAGCAGTAAAGGAGAAGCTGTGATTAAAG | |
| LD2598 | ggaattgtgagcggataacaatttcacacaggaaacagctGATTGCCGACCGCCTGGCCA | pLD3198 |
| LD2599 | GTTCTTCTCCTTTACTCAT ctagtaGTCCTGTGTGA ACGGCGGCGATCAGGGCA | |
| LD2600 | GCTGCCCTGATCGCCGCCGTTCACACAGGACtactagATGAGTAAAGGAGAAGAACTTTT | |
| LD2601 | ACTGCGGTTGGGAAGCTCATGCCGGATCCTTGGCCTGATTTGTATAGTTCATCCATGCc | |
| LD2602 | GGCATGGATGAACTATACAAATCAGGCCAAGGATCCGGCATGAGCTTCCCAACCGCAGT | |
| LD2603 | caaattctgttttatcagaccgcttctgcgttctgatATGATGAGGGCGGTAATGAGGAG | |
| LD3350 | cctgcaggtcgactctagag GCGACGTTTCACAAGACAGA | pLD3791 |
| LD3356 | acagcttctcctttactcatctagtaGTCCTGTGTGACCGGTGGTGGCTACGATT | |
| LD3357 | AATCGTAGCCACCACCGGTCACACAGGACtactagatgagtaaaggagaagctgt | |
| LD3358 | CTTGCCGAAGCCGAACATGCCGGATCCTTGGCCTGAtttgtatagttcatccatgcc | |
| LD3359 | ggcatggatgaactatacaaaTCAGGCCAAGGATCCGGCATGTTCGGCTTCGGCAAG | |
| LD3355 | cagctatgaccatgattacg ATGGATGTGCAGGACCATCT | |
| LD3221 | ggaattgtgagcggataacaatttcacacaggaaacagctGTCCAGGTAGTGGCGAAACA | pLD3685 |
| LD3222 | GCTTCGGCAAGAAATCCGGCAAGGACACCCAATCGTAG | |
| LD3223 | CTACGATTGGGTGTCCTTGCCGGATTTCTTGCCGAAGC | |
| LD3224 | aggcaaattctgttttatcagaccgcttctgcgttctgat CCGCCTCACAGACTGCTC | |

#### Supplementary Table 3: Plasmids

| Plasmids | Number | Description | Source |
|---|---|---|---|
| pMQ30 | LD621 | Yeast-based allelic-exchange vector; *sacB*+, CEN/ ARSH, URA3+, GmR . | (8) |
| pLD3208 | LD3208 | MCS-*mScarlet* GmR, TetR flanked by Flp recombinase target (FRT) sites to resolve out resistance cassettes. Cloned by swapping gfp sequence with mScarlet (XhoI + SacI) | This study |
| pFLP2 | LD743 | Site-specific excision vector with cI857-controlled FLP recombinase encoding sequence, sacB, ApR. | (9) |
| pLD3210 ($P_{rebP1}$-*mScarlet*) | LD3210 | 487 bp of *rebP1* promoter sequence inserted at the MCS (SpeI and EcoRI) of pLD3208 | This study |
| pLD3655 | LD3655 | *gfp* coding region cloned into pAKN69 plasmid replacing the *yfp* coding region excised with SphI and NheI | This study |
| pLD3293 (*mScarlet*+) | LD3293 | *lac*-derived constitutive promoter PA1/04/03 inserted at the MCS (SpeI and EcoRI) of pLD2722 | This study |
| pLD3136 | LD3136 | Δ*PA14_27700 (rcgA)-27690 (fecI2)* flanking fragments introduced into pMQ30 by gap repair cloning in yeast strain InvSc1 | This study |
| pLD3142 | LD3142 | Δ*PA14_27690 (fecI2)* flanking fragments introduced into pMQ30 by gap repair cloning in yeast strain InvSc1 | This study |
| pLD3143 | LD3143 | Δ*PA14_27700 (rcgA)* flanking fragments introduced into pMQ30 by gap repair cloning in yeast strain InvSc1 | This study |
| pLD3179 | LD3179 | Full genomic sequence of *PA14_27700 (rcgA)-27690 (fecI2)* PCR fragment introduced into pMQ30 by gap repair cloning in yeast strain InvSc1. Verified by illumina sequencing. | This study |
| pLD2934 | LD2934 | Δ*rebP1P2* flanking fragments introduced into pMQ30 by gap repair cloning in yeast strain InvSc1 | This study |
| pLD3016 | LD3016 | Full genomic sequence of *rebP1P2* and native promoter PCR fragment introduced into pMQ30 by gap repair cloning in yeast strain InvSc1. Verified by illumina sequencing. | This study |
| pLD3433 | LD3433 | *mScarlet* coding region cloned into pAKN69 plasmid replacing the *yfp* coding region that was excised with BlpI and NheI. | This study |

**Supplementary File 1.** *Pseudomonas aeruginosa* strains that contain homologues of *rebP1* (*PA14_27630*), *fecI2* (*PA14_27690*) and the *reb* gene cluster *PA14_27630-PA14_27700*. Strains were identified with a nucleotide NCBI BLAST against 312 complete genomes in *Pseudomonas aeruginosa* (taxid:287). 189 strains contained the *reb* cluster with *rebP1* and *fecI2*; in one strain the cluster lacked *fecI2*. 122 strains lacked the *reb* cluster, *rebP1* and *fecI2*.

**Supplementary File 2.** List of all 126 proteins (represented by two or more peptides) found in the SDS-insoluble fraction by tandem mass spectrometry.

#### SUPPLEMENTARY REFERENCES


1. Raymann, K., Bobay, L.-M., Doak, T. G., Lynch, M. & Gribaldo, S. A genomic survey of Reb homologs suggests widespread occurrence of R-bodies in proteobacteria. *G3* **3**, 505–516 (2013).

2. Rahme, L. G. *et al.* Common virulence factors for bacterial pathogenicity in plants and animals. *Science* **268**, 1899–1902 (1995).

3. Madsen, J. S. *et al.* Facultative control of matrix production optimizes competitive fitness in *Pseudomonas aeruginosa* PA14 biofilm models. *Appl. Environ. Microbiol.* **81**, 8414–8426 (2015).

4. Liberati, N. T. *et al.* An ordered, nonredundant library of *Pseudomonas aeruginosa* strain PA14 transposon insertion mutants. *Proc. Natl. Acad. Sci. U. S. A.* **103**, 2833–2838 (2006).

5. Brenner, S. The genetics of *Caenorhabditis* elegans. *Genetics* **77**, 71–94 (1974).

6. Simon, R., Priefer, U. & Pühler, A. A Broad Host Range Mobilization System for In Vivo Genetic Engineering: Transposon Mutagenesis in Gram Negative Bacteria. *Biotechnology* **1**, 784–791 (1983).

7. Dehio, C. & Meyer, M. Maintenance of broad-host-range incompatibility group P and group Q plasmids and transposition of Tn5 in Bartonella henselae following conjugal plasmid transfer from *Escherichia coli*. *J. Bacteriol.* **179**, 538–540 (1997).

8. Shanks, R. M. Q., Caiazza, N. C., Hinsa, S. M., Toutain, C. M. & O'Toole, G. A. *Saccharomyces cerevisiae*-based molecular tool kit for manipulation of genes from gram-negative bacteria. *Appl. Environ. Microbiol.* **72**, 5027–5036 (2006).

9. Hoang, T. T., Karkhoff-Schweizer, R. R., Kutchma, A. J. & Schweizer, H. P. A broad-host-range Flp-FRT recombination system for site-specific excision of chromosomally-located DNA sequences: application for isolation of unmarked *Pseudomonas aeruginosa* mutants. *Gene* **212**, 77–86 (1998).
